# A PKA Inhibitor Motif within Smoothened Controls Hedgehog Signal Transduction

**DOI:** 10.1101/2021.07.05.451193

**Authors:** John T. Happ, Corvin D. Arveseth, Jessica Bruystens, Daniela Bertinetti, Isaac B. Nelson, Cristina Olivieri, Danielle S. Hedeen, Ju-Fen Zhu, Jacob L. Capener, Jan Wilfried Bröckel, Lily Vu, C.C. King, Victor L. Ruiz-Perez, Gianluigi Veglia, Friedrich W. Herberg, Susan S. Taylor, Benjamin R. Myers

## Abstract

The Hedgehog (Hh) cascade is central to development, tissue homeostasis, and cancer. A pivotal step in Hh signal transduction is the activation of GLI transcription factors by the atypical G protein-coupled receptor (GPCR) Smoothened (SMO). How SMO activates GLI has remained unclear for decades. Here we show that SMO employs a decoy substrate sequence to physically block the active site of the PKA catalytic subunit (PKA-C) and extinguish its enzymatic activity. As a result, GLI is released from phosphorylation-induced inhibition. Using a combination of *in vitro*, cellular, and organismal models, we demonstrate that interfering with SMO / PKA pseudosubstrate interactions prevents Hh signal transduction. The mechanism we uncovered echoes one utilized by the Wnt cascade, revealing an unexpected similarity in how these two essential developmental and cancer pathways signal intracellularly. More broadly, our findings define a new mode of GPCR-PKA communication that may be harnessed by a range of membrane receptors and kinases.

## INTRODUCTION

The Hh signaling cascade is fundamental to embryogenesis, controlling the development of nearly every vertebrate organ^1-4^. Insufficient Hh pathway activity underlies birth defects affecting the nervous, cardiovascular, and musculoskeletal systems^5-7^. On the other hand, Hh pathway overactivation drives several common cancers, including basal cell carcinoma of the skin (the most common cancer in North America) and medulloblastoma (the most common pediatric brain tumor)^8,9^. The Hh pathway utilizes an unusual signal transduction mechanism involving layers of repressive interactions^1-4^. In the pathway “off” state, PKA-C phosphorylates GLI, stimulating its proteolysis into a truncated transcriptional repressor that inhibits target gene expression^4,10^. In the pathway “on” state, Hh ligands bind to and inactivate the 12-transmembrane sterol transporter PATCHED1 (PTCH1), which releases SMO from PTCH1-mediated inhibition^4^. This process allows SMO to access its endogenous sterol ligands and undergo an activating conformational change^11-14^. Once activated, SMO blocks PKA-C-mediated phosphorylation of GLI^15-18^, a key step that likely occurs within the tiny cell-surface compartment formed by the primary cilium^19-21^. Consequently, GLI is activated and can control the expression of proliferative or differentiative genes^10^. SMO regulation of PKA-C is thus a critical event in transducing Hh signals from the cell surface to the nucleus. However, the underlying mechanism has remained mysterious for decades^1-4^.

GPCRs classically inhibit PKA-C via well-characterized signaling cascades involving heterotrimeric G protein-mediated effects on cAMP which promote formation of inactive PKA holoenzymes^22,23^. In contrast, we recently found that SMO prevents PKA-C from phosphorylating substrates via a noncanonical mechanism. SMO directly interacts with PKA-C subunits, recruiting them to the membrane and thereby restricting their access to soluble GLI proteins. SMO / PKA-C interactions are triggered by GPCR kinases 2 and 3 (GRK2/3), which recognize the SMO active state and phosphorylate the cytoplasmic tail (C-tail) of SMO to promote PKA-C binding. Based on these observations, we proposed that active, phosphorylated SMO binds to and sequesters PKA-C within the cilium, which prevents phosphorylation of GLI and thereby promotes GLI activation^24^.

Here we uncover a critical and unexpected component of the SMO / PKA-C regulatory mechanism in which SMO physically blocks PKA-C enzymatic activity. We show that the SMO proximal C-tail (pCT) acts as a decoy PKA-C substrate that binds to and occludes the kinase active site, thereby preventing phosphorylation of PKA-C targets. SMO is, to our knowledge, the first example of a GPCR that functions as a direct PKA-C inhibitor. However, this decoy substrate mechanism appears to apply more generally to transmembrane receptors and kinases in other signaling pathways.

## RESULTS

### SMO binds and inhibits PKA-C as a pseudosubstrate

The principal regulators of PKA-C activity within cells are PKA regulatory (PKA-R) subunits^25^ and the heat-stable protein kinase inhibitor (PKI) proteins^26^. PKI proteins and type I (PKA-RI) regulatory subunits operate as pseudosubstrates that bind within the PKA-C active site but cannot undergo phosphorylation. As a result, the enzyme’s phosphoryl transfer and substrate turnover cycle is interrupted^25-28^. PKA-R / PKA-C holoenzymes dissociate upon cAMP binding to PKA-R, releasing catalytically active PKA-C^25^. In contrast, PKI proteins interact with PKA-C independently of cAMP levels^26^. Despite their divergent regulatory influences, PKI and PKA-RI engage the PKA-C active site cleft using similar sequence motifs^25,28^.

Inspection of the mouse SMO sequence revealed a region within the pCT (residues 615-638) that bears the hallmarks of a PKA-C pseudosubstrate (Fig. 1a). First, in contrast to canonical PKA-C substrates, which contain a serine or threonine at a canonical phosphorylation site (P-site)^29^, SMO possesses a non-phosphorylatable residue, alanine, at this position. Second, SMO contains arginines at the P-2 and P-3 positions that are essential in other pseudosubstrates (PKI and PKA-RI) for binding the PKA-C active site^30-33^. Third, SMO contains a hydrophobic residue, isoleucine, at P+1 and a bulkier aromatic residue, tryptophan, at P-13, both of which contribute to high-affinity PKI interactions with PKA-C^34-36^. Finally, SMO harbors a predicted α-helical sequence N-terminal to the pseudosubstrate region (Fig. 1a), resembling a domain in PKI that is required for high-affinity interactions with PKA-C^37,38^. Based on these observations and the studies described below, we refer to SMO residues 615-638 as the “SMO PKI motif”.

**Fig. 1:**
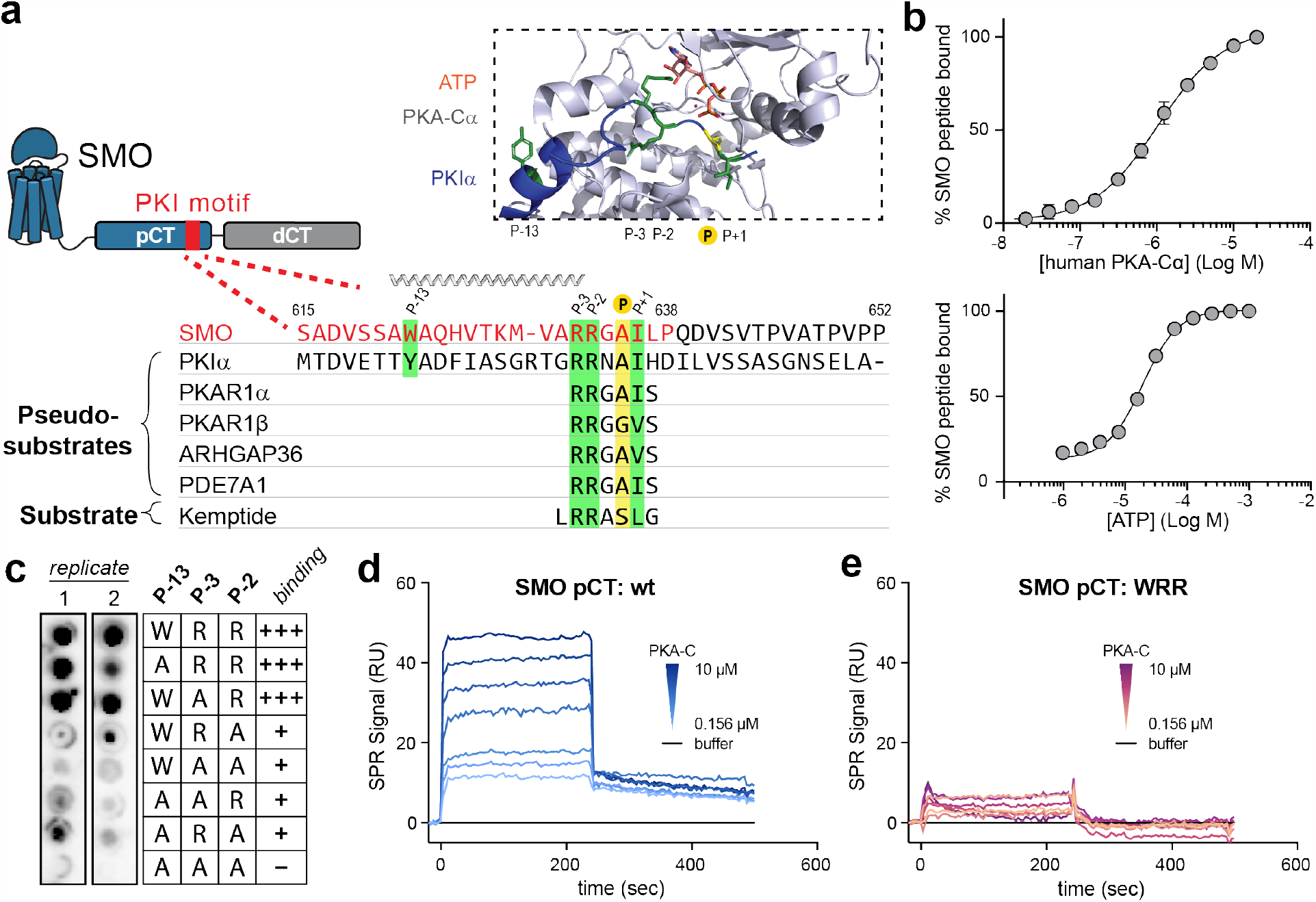
SMO binds PKA-C as a pseudosubstrate. **a**, CLUSTAL alignment of the mouse SMO pCT with the PKIɑ pseudosubstrate region. Additional PKA-C pseudosubstrate and substrate sequences are provided for comparison^27,96,97^. P-site is yellow; other key conserved residues are green. Spiral cartoon above alignment indicates predicted SMO helical region. Standard (615-638) and extended (615-652) SMO peptides used for *in vitro* assays are colored red or black, respectively. Inset, structure of PKA-Cɑ bound to PKIɑ(5-24) (PDB: 3FJQ), with ATP colored orange and key PKI residues colored as described above. **b**, Top, fluorescence polarization assay employing FAM-labeled SMO peptide, 1 mM ATP, and varying concentrations of human PKA-Cɑ. Bottom, the same assay except with 3 µM PKA-Cɑ and varying concentrations of ATP. **c**, Overlay of purified mouse PKA-Cɑ onto an array of SMO peptides containing the indicated substitutions in the P-13, P-3, and P-2 positions. **d**, SPR sensorgram for binding of GST-tagged wild-type SMO pCT to a series of PKA-Cɑ concentrations ranging from 0.156 μM to 10 μM. Buffer contained 1 mM ATP and 10 mM MgCl_2_. **e**, As **d**, but with SMO pCT harboring the WRR mutation.

We hypothesized that SMO utilizes its PKI motif as a pseudosubstrate to bind and inhibit PKA-C, thereby activating GLI. Consistent with this hypothesis, residues 615-637 of mouse SMO are essential for communication with GLI^39^, although the structure, function, and interacting partner(s) of this SMO region are all undefined. Furthermore, the P, P+1, P-2, P-3, and P-13 residues critical for PKA-C pseudosubstrate function in *bona fide* PKI proteins are highly conserved among SMO orthologs (Extended Data Fig. 1), suggestive of a central role in SMO functionality.

We measured the affinity of the SMO PKI motif for PKA-Cα, the best-studied and most ubiquitously expressed PKA-C isoform^27^, using fluorescence polarization assays. A fluorescently labeled peptide encompassing the SMO PKI motif (standard SMO peptide, Fig. 1a, red) bound saturably to purified human (K_D_ = 823 nM, Fig. 1b (top)) or mouse (K_D_ = 911 nM, Extended Data Fig. 2a (left)) PKA-Cα. These interactions showed pseudosubstrate characteristics: they were strongly ATP-dependent^40^ (EC_50_ for ATP = 19.2 µM (human) or 0.59 µM (mouse), Fig. 1b (bottom) and Extended Data Fig. 2a) (right)) and were blocked by alanine substitution of the P-2 / P-3 arginines and the P-13 tryptophan, hereafter referred to as the “WRR mutation” (Fig. 1c, Extended Data Fig. 2b). Along similar lines, direct surface plasmon resonance (SPR) binding studies revealed that a recombinant protein encompassing the entire 93 amino-acid SMO pCT bound specifically to PKA-Cα (Fig. 1d). These interactions displayed fast on- and off-rates with transient kinetics, and steady-state analysis of the binding data revealed a K_D_ of 835 nM (Extended Data Fig. 2c,d). Consistent with our peptide binding studies, PKA-C interactions with the SMO pCT depended strictly on ATP and MgCl_2_ (Extended Data Fig. 2e), and only minimal PKA-C binding was observed for the WRR mutant (Fig. 1e, Extended Data Fig. 2c,d). Interestingly, the above SMO PKI binding assays revealed weaker interactions with PKA-C than observed for conventional pseudosubstrates (K_D_ = 1.1 nM for PKIα(5-24) peptide^41^). We discuss the biochemical basis and biological significance for these affinity differences below.

We next studied the interaction between PKA-C and the SMO PKI motif at a structural level using nuclear magnetic resonance (NMR) spectroscopy. Binding of pseudosubstrate sequences from either SMO or PKIɑ induced similar overall chemical shift perturbations throughout the PKA-C kinase core (Fig. 2a, Extended Data Fig. 2f). SMO PKI peptides also blocked PKA-C enzymatic activity *in vitro*, as assessed via a PKA-C substrate phosphorylation assay^42^, with an extended length peptide (Fig. 1a) inhibiting PKA-C more profoundly (Fig. 2b). In keeping with this trend, the recombinant SMO pCT (Extended Data Fig. 2g,h) required 5-10-fold lower concentrations to efficiently block PKA-C activity compared to SMO PKI peptides (24-38 amino acids in length) (IC_50_ = 10.9 µM for pCT vs. 50-125 µM for SMO PKI peptides, Fig. 2c). Consistent with a pseudosubstrate mode of inhibition, introducing the WRR mutation into the SMO pCT restored PKA-C activity (Fig. 2d). These data indicate that the SMO PKI motif is sufficient to inhibit PKA-C substrate phosphorylation, with additional sequences in the pCT enhancing the efficiency of inhibition.

**Fig. 2:**
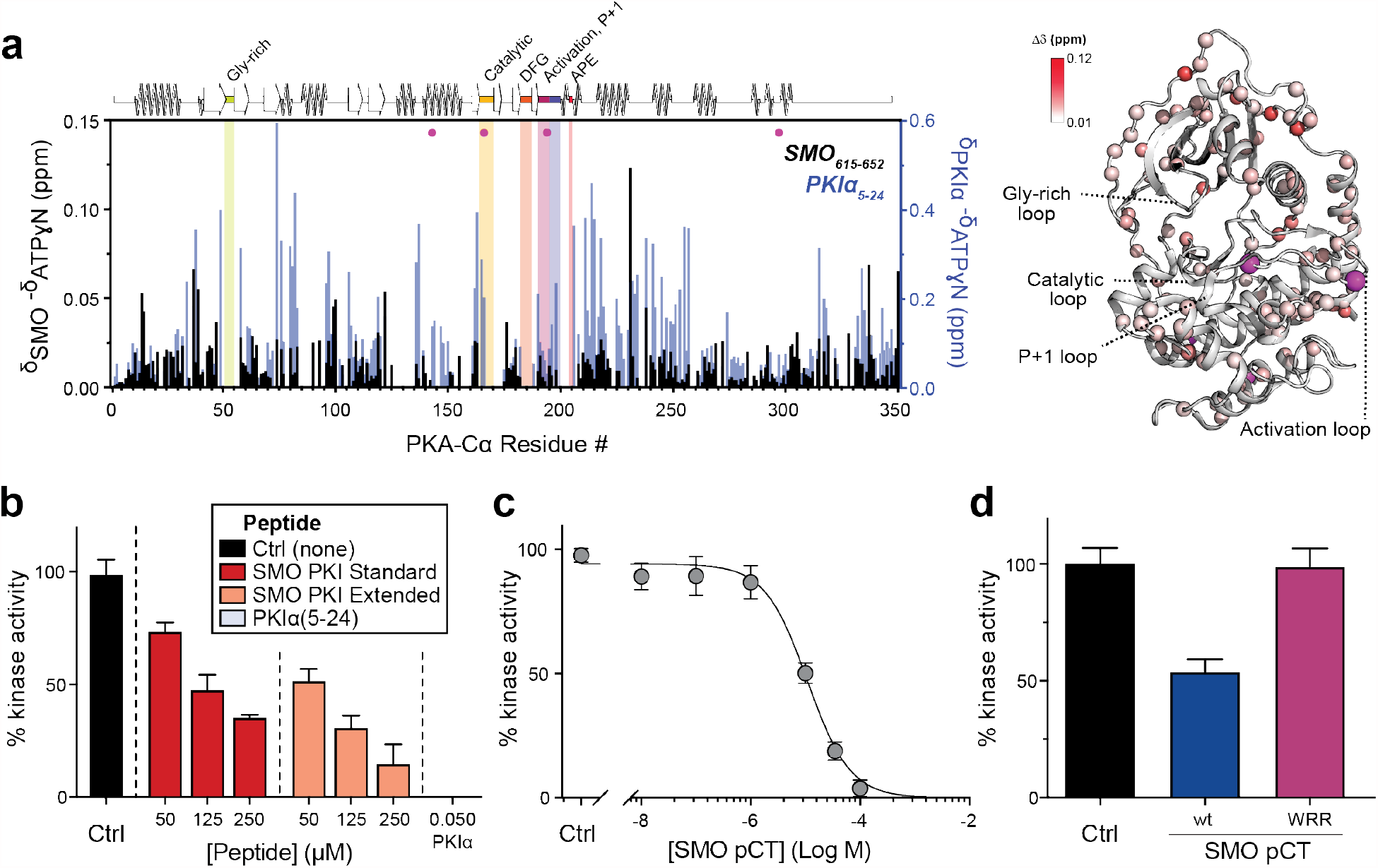
SMO is a pseudosubstrate inhibitor of PKA-C. **a**, Mapping of amide backbone chemical shift perturbations (CSP, δ) for [^1^H, ^15^N]-labeled PKA-Cɑ bound to nucleotide (ATPɑN) and either an extended SMO PKI peptide (black) or a control PKIɑ(5-24) peptide^90^ (blue), calculated relative to ATPγN-bound PKA-Cɑ without peptide. Key functional domains of PKA-C are highlighted along the X-axis. Magenta spheres indicate PKA-Cɑ residues (R144, D166, R194, and F297) that show a signal for PKIɑ(5-24) but not SMO. Right, CSP values were mapped onto the PKA-Cɑ structure (PDB: 4WB5) and displayed as a heatmap. **b**, Spectrophotometric assay of PKA-Cɑ substrate phosphorylation, in the presence of standard (red) or extended (coral) SMO peptides (see Fig. 1a) or a control PKIα(5-24) peptide (grey) **c**, Concentration dependent inhibition of PKA-Cɑ with recombinant SMO pCT. **d**, As **b**, but comparing wild-type vs. WRR mutant versions of the recombinant SMO pCT. Inhibition in **b-d** is calculated relative to a control without SMO peptide (Ctrl.). See Extended Data Table 1 for statistical analysis.

### The SMO PKI motif is required for Hh signal transduction

We next asked whether the SMO PKI motif contributes to PKA-C binding and inhibition in living systems. Our initial studies involved a simplified HEK293 cell model for SMO / PKA-C regulation and employed truncated SMO constructs (either SMO657 or SMO674, with the number indicating the C-terminal-most residue) that exhibit improved expression and biochemical stability compared to full-length SMO^24^. We used bioluminescence resonance energy transfer (BRET)^43,44^ to detect interactions between nanoluciferase (nanoluc)-tagged SMO and YFP-tagged PKA-C constructs in a cellular environment^24^. In this assay, PKA-C exhibited strong, specific BRET with wild-type SMO657, but not with SMO657 harboring the WRR mutation (Fig. 3a). Coimmunoprecipitation assays confirmed these results (Extended data Fig. 3a). Similarly, the WRR mutation prevented SMO674 from colocalizing with PKA-C at the membrane^24^, as determined by live-cell confocal microscopy (Fig. 3b).

**Fig. 3:**
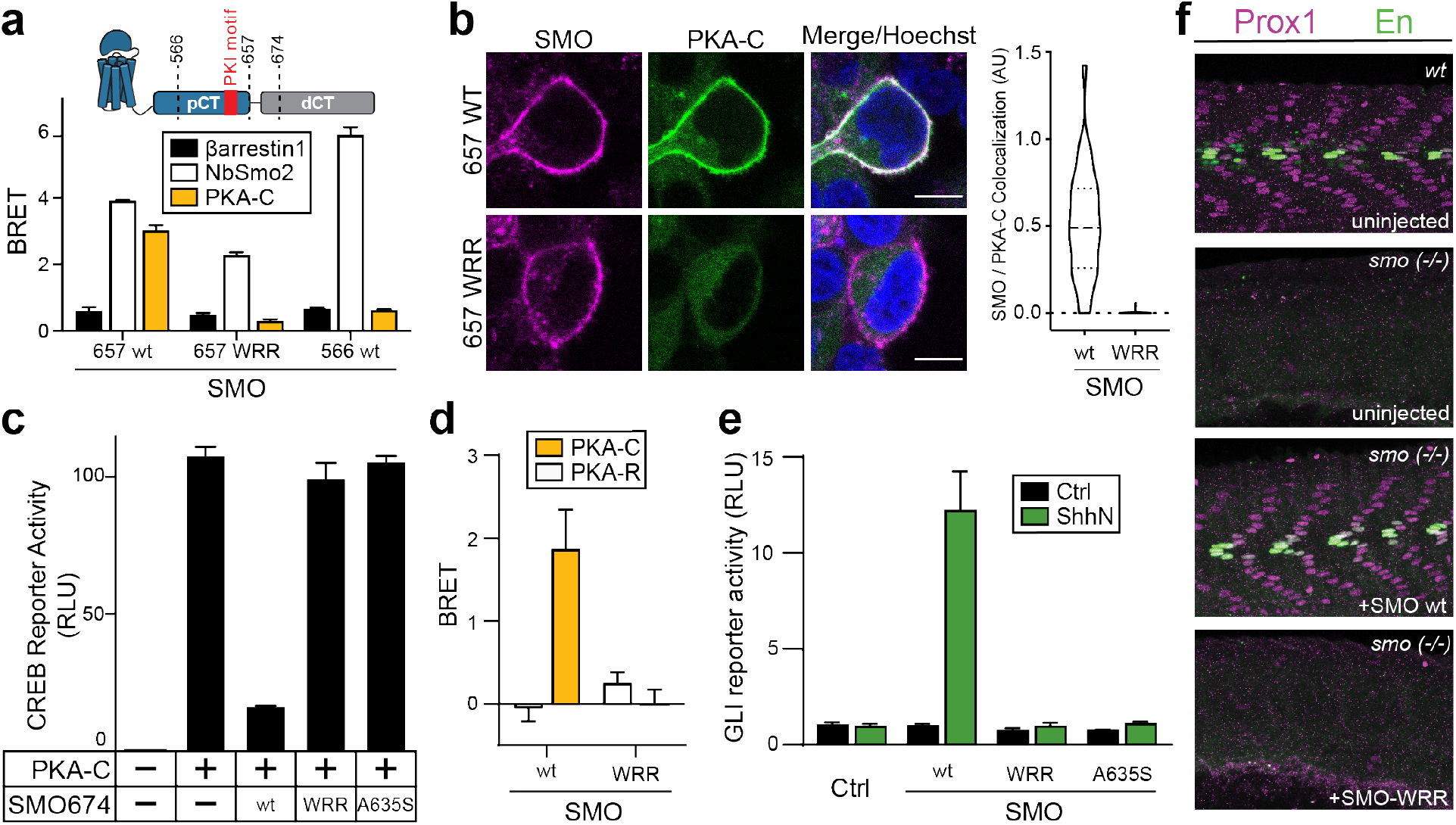
The SMO PKI motif is required for Hh signal transduction. **a**, Top, schematic diagram of truncated SMO expression constructs. Bottom, BRET analysis of SMO / PKA-C interactions in HEK293 cells expressing nanoluc-tagged wild-type (wt) SMO657 or SMO657 harboring the WRR mutation (WRR), along with YFP-tagged PKA-Cɑ. SMO566 (which lacks the C-tail) serves as a negative control donor. YFP-tagged βarrestin1 (which exhibits minimal binding to SMO) and NbSmo2 (which binds the intracellular surface of the SMO 7TM domain) serve as negative and positive control acceptors, respectively^24^. **b**, Left, confocal images of live HEK293 cells coexpressing GFP-tagged PKA-Cɑ with FLAG-tagged wild-type or mutant SMO674. Cells were treated with SMO agonist (SAG21k). Scale bar = 10 µm. Right, quantification of SMO / PKA-C colocalization (n=34-48 cells per condition). **c**, CREB transcriptional reporter assay, reflecting PKA-C mediated substrate phosphorylation, in HEK293 cells transfected with PKA-Cɑ and the indicated SMO674 constructs. **d**, BRET analysis of SMO / PKA-C interactions in IMCD3 cells expressing nanoluc-tagged wild-type or mutant SMO, along with low levels of PKA-Cɑ-YFP. PKA-RIɑ-YFP serves as a negative control. Under these conditions, PKA-Cɑ-YFP is expressed at substantially lower levels than PKA-RIɑ-YFP^24^. **e**, GLI transcriptional reporter assay in *Smo*^*-/-*^ MEFs transfected with a GFP negative control (Ctrl.), or the indicated wild-type (wt) or mutant SMO constructs. Cells were treated with conditioned medium containing the N-terminal signaling domain of Sonic hedgehog (ShhN), or control, non-ShhN-containing conditioned medium (Ctrl). **f**, Wild-type or *smo*^*-/-*^ zebrafish injected with the indicated mRNA constructs were stained for Prox1 (magenta) or Engrailed (En, green) to mark muscle fiber nuclei. n=12 (uninjected), n=41 (SMO wt), n= 47 (SMO WRR). See Extended Data Table 1 for statistical analysis.

We next assessed the impact of the SMO PKI motif on PKA-C activity in cells using a model PKA-C substrate, the cyclic AMP response element binding protein (CREB) transcription factor^45^, and a CREB transcriptional reporter assay. This experimental paradigm entails overexpression of PKA-C at levels exceeding those of endogenous PKA-R, thereby minimizing potentially confounding contributions from heterotrimeric G protein- and cAMP-containing cascades^24^. Under these conditions, wild type SMO674 blocked PKA-C-mediated CREB reporter activation while SMO674 harboring the WRR mutation failed to do so (Fig. 3c). We observed similar effects with serine substitution of the P-site alanine (A635S, Fig. 3c and Extended data Fig. 3a), which converts pseudosubstrates to substrates and thereby promotes their dissociation from the PKA-C active site^33,46^.

We extended the above findings to more physiological cellular models of Hh signaling in primary cilia, using SMO constructs with full-length unmodified C-termini. We studied SMO / PKA-C colocalization in cilia using inner medullary collecting duct (IMCD3) cells, a robustly ciliated kidney cell line that is used extensively in studies of ciliary Hh signal transduction^24,47-49^. Upon Hh pathway activation, SMO accumulates in IMCD3 primary cilia^47-49^, where it colocalizes with PKA-C^24^. In contrast, PKA-C displayed dramatically reduced colocalization in cilia with SMO containing the WRR mutation (Extended Data Fig. 3b). The WRR mutation also prevented interaction of SMO with PKA-C (expressed at near-endogenous levels^24^) in these ciliated cells (Fig. 3d).

To capture the entire process of Hh signal transduction from the cell surface to the nucleus, we used mouse embryonic fibroblasts (MEFs) and a GLI transcriptional reporter assay^11,50,51^. This model strictly requires SMO, PKA-C, GLI, and an intact cilium^50,52,53^. In *Smo*^*-/-*^ MEFs, transfection of wild-type SMO enabled strong GLI transcriptional responses to Hh ligands, whereas the WRR and A635S mutants were almost completely devoid of activity (Fig. 3e).

Mutations in the SMO PKI motif specifically affect binding to and inhibition of PKA-C, based on the following control studies. First, the expression levels and electrophoretic mobilities of these mutants were similar to those of their wild-type counterparts (Extended Data Fig. 3a and 4a,b,d), and the mutants localized correctly to primary cilia (Extended Data Fig. 3b, 4c). In addition, the SMO mutations only minimally affected BRET with nanobody 2 (NbSmo2, Fig. 3a), which selectively binds the active state of SMO via the cytoplasmic face of its seven-transmembrane (7TM) domain^24^. Finally, the mutants underwent normal SMO activity-dependent phosphorylation by GRK2/3 kinases^24,54^ (Extended Data Fig. 4d). These data rule out possible nonspecific effects of the mutations on SMO expression, trafficking, conformational activation, or ability to serve as a GRK2/3 substrate.

To examine the role of the SMO PKI motif in Hh signal transduction *in vivo*, we studied the specification of slow muscle cell types in zebrafish embryos, a widely utilized model of morphogenetic Hh signal transduction during vertebrate development^24,55,56^. While expression of wild-type SMO restored correct muscle specification to *smo*^*-/-*^ zebrafish, the WRR or A635S mutants did not (Fig. 3f, Extended Data Table 2).

Taken together, our studies demonstrate a requirement for PKA-C pseudosubstrate interactions in SMO inhibition of PKA-C, and activation of GLI, during Hh signal transduction in both cultured cells and organisms.

### An avidity-based mechanism for SMO inhibition of PKA-C

PKA-C interactions with the SMO PKI motif, while essential for Hh signal transduction (Fig. 3), appear weaker (K_D_ = 823-911 nM for SMO PKI peptides, Fig. 1b, Extended Data Fig. 2a,d) than those with a canonical PKA-C pseudosubstrate, (K_D_ = 1.1 nM for PKIα(5-24) peptide^41^). SMO / PKA-C interactions, however, are also influenced by sequences outside the PKI motif, indicating that they are more favorable *in vivo* than the *in vitro* measurements suggest. Several lines of evidence support this proposal. First, SMO mutations known to reduce PKA-C interactions *in vivo*, including deletion of a predicted amphipathic helix spanning residues 570-581 or alanine substitution of GRK2/3 phosphorylation sites^24^, map to non-PKI regions of the pCT (summarized in Fig. 4a). Second, SMO pCT interactions with PKA-C are enhanced by the distal SMO C-tail (dCT, residues 658-793)^24^, demonstrating that this region harbors additional PKA-C binding determinants. Accordingly, in HEK293 cells, PKA-C interactions with SMO containing its full-length C-tail are only partially blocked by the WRR mutation (Fig. 4b), in contrast to the near-complete loss of interaction with SMO lacking the dCT (Fig. 3a), Finally, a tiled SMO C-tail peptide microarray identified several PKA-C-binding sequences in addition to the PKI motif, including one in the dCT with PKI-like attributes (Fig. 4c, Extended Data Fig. 5).

**Fig. 4:**
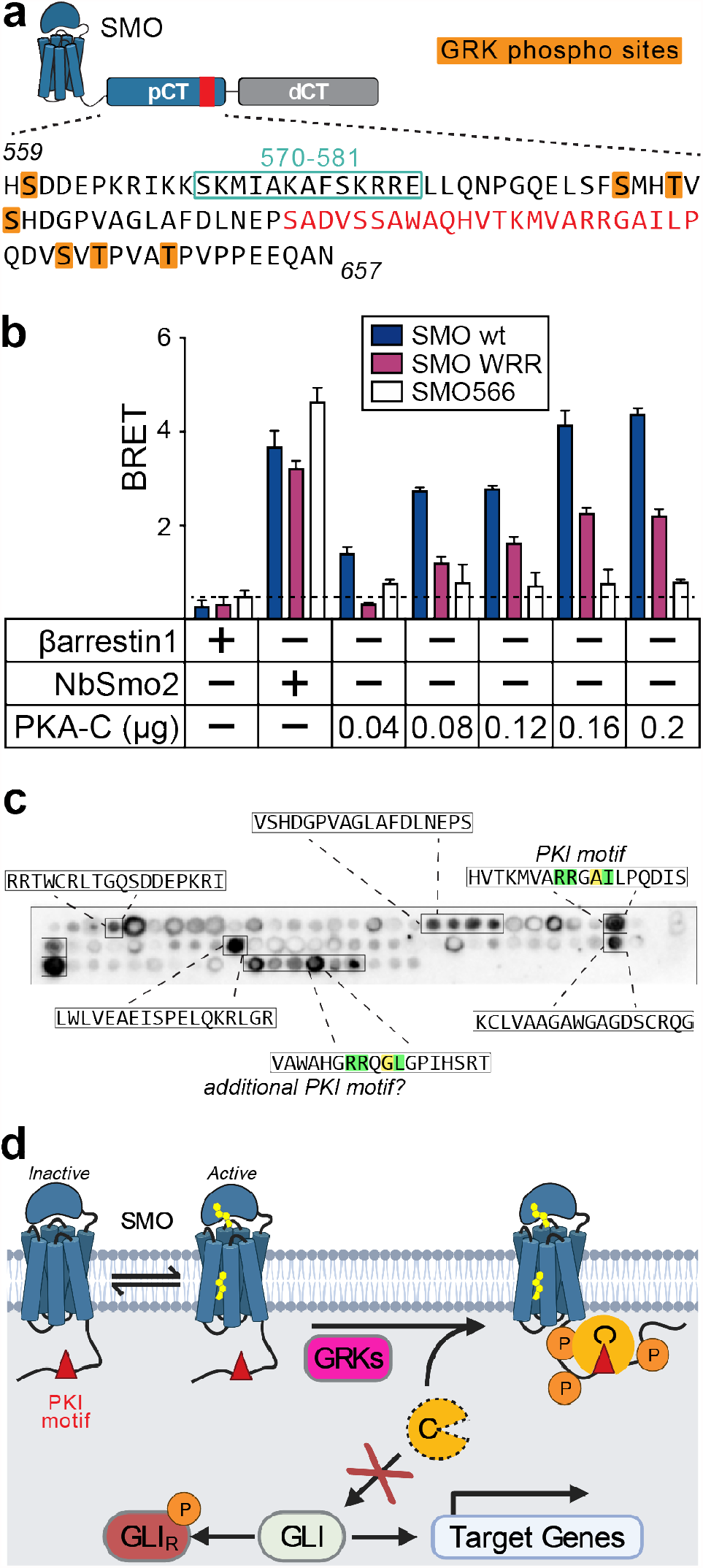
An avidity-based mechanism for SMO inhibition of PKA-C. **a**, Annotated sequence of the mouse SMO pCT. PKI motif is indicated in red, along with GRK2/3 phosphorylation sites (orange) and residues 570-581 (aquamarine box), previously shown to influence SMO / PKA-C interactions and Hh signal transduction^24,61^. **b**, BRET analysis of SMO / PKA-C interactions in HEK293 cells transfected with full-length wild-type SMO (wt, navy) WRR mutant (purple), or C-terminally truncated (SMO566, white) versions of SMO as donor, and the indicated DNA amounts of PKA-Cɑ as acceptor. Nonspecific signal is indicated by the negative control BRET acceptor βarrestin1 (dashed line). See Extended Data Table 1 for statistical analysis. **c**, Representative image of a tiled array of 18mer peptides covering the complete C-tail of human SMO, probed with PKA-Cɑ as in Fig. 1c. Peptides that bind are boxed and their sequences indicated. **d**, Proposed model for Hh signal transduction, as described in “Discussion”.

Taken together, our data support an avidity-based mechanism for SMO / PKA-C binding. A set of ancillary interactions cooperate to stabilize the core binding events between the SMO PKI motif and the PKA-C active site. This avidity-based mechanism may provide the essential link between SMO activation and disruption of PKA-C phosphoryl transfer during Hh signal transduction.

## DISCUSSION

SMO inhibition of PKA-C is a central aspect of Hh signal transduction in development and disease. Instead of obeying the existing paradigms for GPCR-PKA signaling, SMO enjoys a private signaling pathway whereby it directly binds PKA-C as a pseudosubstrate. This both restricts access of PKA-C to soluble targets and extinguishes its enzymatic activity. A PKI motif encoded by the SMO cytoplasmic domain is central to this mechanism in two ways. First, the PKI motif sequesters PKA-C away from GLI by promoting SMO / PKA-C interactions, and thus recruitment of PKA-C to the membrane. Second, the PKI motif occludes the PKA-C active site, interrupting catalysis. By utilizing this two-pronged strategy, SMO can efficiently block PKA-C-mediated phosphorylation, activate GLI, and promote transcription of Hh pathway target genes.

Based on our findings, we propose a revised model for Hh signal transduction downstream of SMO (Fig. 4d). In the pathway “off” state, SMO is in an inactive conformation^11,51,57,58^. As a result, SMO cannot be phosphorylated efficiently by GRK2/3^24,54^, and exists at low levels in the ciliary membrane^59-61^. We expect interactions between PKA-C and the SMO PKI motif to be unfavorable under these conditions due to their distinct subcellular localizations and relatively weak affinity. In the pathway “on” state, Hh-mediated inactivation of PTCH1 enables SMO to bind its endogenous sterol ligands and adopt an active conformation^11-13,51^. Consequently, SMO undergoes phosphorylation by GRK2/3 and accumulates in the cilium, bringing SMO in close proximity to ciliary PKA-C. This allows SMO to bind PKA-C via its PKI motif, which both sequesters PKA-C at the ciliary membrane^24^ and blocks PKA-C-mediated catalysis. These interactions are facilitated by other PKA-C-binding sequences in the SMO pCT and dCT. We expect the interactions are also enhanced by the membrane itself, which restricts diffusion of proteins to two dimensions and can thereby dramatically increase effective concentrations relative to soluble compartments^62^. Additionally, SMO activation could conceivably decrease ciliary cAMP via coupling to inhibitory (Gα_i/o/z_) G protein cascades^51,63,64^ or by triggering the ciliary exit of GPR161, a constitutively active GPCR that couples to stimulatory (Gα_s_) G proteins^47,49,65^. These processes, while not absolutely required for Hh signal transduction^65-70^, may reduce levels of free (non-PKA-R-bound) PKA-C in cilia^71^, and thereby aid the SMO PKI motif in binding and inhibiting the ciliary pool of PKA-C. Thus, activation of SMO prevents PKA-C substrate phosphorylation via a host of auxiliary interactions and regulatory influences that assist the essential pseudosubstrate action of the SMO PKI motif.

Our study helps explain a crucial but enigmatic aspect of Hh signal transduction, namely how PKA-C is inhibited, and GLI activated, only when the Hh pathway is in an “on” state. Indeed, our findings are consistent with a number of prior observations, such as that inactivation of GRK2/3 decreases SMO / PKA-C interactions^24^, blocks SMO / PKA-C colocalization at the membrane^24^, and abolishes Hh signal transduction^56,72,73^. Based on our model, these effects can be explained by GRK2/3 inhibition weakening the SMO / PKA-C interaction below a critical threshold such that the PKI motif cannot efficiently engage the enzyme’s active site. Furthermore, our findings suggest that SMO / PKA-C interactions are appropriately tuned: If the SMO PKI motif were to bind PKA-C with high affinity, the Hh pathway could misfire, as PKA-C would be inhibited (and GLI dephosphorylated) even when SMO is inactive. Conversely, reducing the affinity of PKA-C for the SMO PKI motif, as occurs with the SMO WRR or A635S mutations, prevents the SMO active state from inhibiting PKA-C and thereby hinders GLI transcription. Future structural studies of full-length SMO / PKA-C complexes will provide a more detailed understanding of how the SMO pCT engages PKA-C, and may also reveal precisely how GRK2/3 facilitates these interactions.

The mechanism described above represents, to our knowledge, an unprecedented mode of GPCR-kinase communication. It is, however, remarkably similar to a strategy used by another critical developmental and cancer pathway: the Wnt cascade^74-77^ (Extended Data Fig. 6). During the initial steps of Wnt signal transduction at the plasma membrane, the cytoplasmic tail of LDL receptor-related proteins 5/6 (LRP5/6) serves as a pseudosubstrate inhibitor of GSK-3β kinases. This blocks phosphorylation and degradation of the transcriptional coactivator β-catenin^78-81^. Moreover, the LRP5/6 tail binds GSK-3β with low affinity, but Wnt pathway activation enhances this interaction through phosphorylation of the LRP5/6 tail as well as recruitment of GSK-3β-containing protein complexes to the membrane^78-80^. Thus, in both the Hh and Wnt cascades, a phosphorylated receptor tail sequesters and competitively inhibits a kinase via an avidity-based mechanism. It was unexpected that these two pathways, which utilize distinct molecular components and rely on disparate subcellular environments (cilium vs. plasma membrane)^21,82^, nevertheless share a common mechanism for intracellular signal transduction. Why such a mechanism evolved to carry out central steps in these pathways is presently unclear. One possibility is that it serves to spatially restrict the kinase-inhibiting effects of upstream receptors and thereby avoid pleiotropic effects of inhibiting these kinases globally. This may help to ensure signaling specificity and reduce untoward crosstalk with other pathways^75,78,81^. It is also possible that such a mechanism is particularly well-suited to the embryogenesis- and homeostasis-related functions of Hh and Wnt signaling. In any event, the use of a similar strategy during Hh and Wnt signal transduction hints at a more general applicability to a range of transmembrane receptors and kinases. Given the ubiquity of these proteins in metazoan physiology and disease^22,23,28^, and their prevalence as therapeutic targets^83,84^, this represents an exciting area for future study.

## METHODS

### Cell culture and zebrafish husbandry

HEK293FT cells, IMCD3 Flp-in cells, *Smo*^-/-^ MEFs, and HEK293-Freestyle cells were grown as previously described^24^. Stably transfected HEK293 Flp-in T-rex cells were constructed and maintained as previously described^11^. IMCD3 stable line expressing SMO(WRR)-IRES-PKACmNG produced as previously described, others (PKAC-mNG and 2ARNb80-mNG) already described. Zebrafish were maintained as previously described^24^.

### Antibodies, small molecules, and other reagents

AG21k was a gift from P. Beachy. Cmpd101 was obtained from Hello Bio. Rabbit anti-GFP (which also detects YFP) was obtained from Thermo Fisher Scientific (A11122). Mouse anti-Arl13b was obtained from Antibodies Inc (75-287). Alexa 647-conjugated M1 FLAG antibody and M1 FLAG affinity resin were prepared in-house. Control or ShhN conditioned medium was prepared from stably transfected HEK293 cells as previously described^24^. Dual-luciferase assay was obtained from Promega. Coelenterazine h was obtained from NanoLight Technology (301–500). Furimazine was obtained from AOBIOUS (AOB36539). For transfection studies, TransIT 2020 and TransIT 293 were obtained from Mirus Bio.

### DNA constructs

For expression and purification of the SMO pCT in *E. coli*, the ShhN gene was excised from ShhN / pHTSHP^51^ and replaced with residues 565-657 of mouse SMO (^565^KRIKK…PEEQAN^657^), downstream of the MBP-His_8_-SUMO tags in the vector pHTSHP, or for GST-constructs in the vector pGexKG. GST-PKI construct in pGexKG was previously described^85^. The resulting construct was termed SMO pCT(565-657). The SMO655-nanoluc, SMO674, and SMO-nanoluc constructs (all in pVLAD6, with N-terminal hemagglutinin signal sequence and FLAG tag) were previously described^24^. Full-length SMO (in pGEN, with C-terminal myc tag) was previously described^86^. Mouse PKA-Cα-YFP, PKA-RIα-YFP, and NbSmo2-YFP constructs in pVLAD6 were previously described^24^. Barrestin1-YFP in pCDNA3.1-zeo was previously described^24^. To construct IMCD3 Flp-in stable lines coexpressing FLAG-tagged mutant SMO with mNeonGreen-tagged PKA-Cα (using an IRES element), the previously described construct SMO-IRES-PKACαmNG / pEF5-FRT^24^ was modified to introduce the WRR mutation into SMO. To construct stable Flp-in HEK293 cell lines coexpressing wild-type or mutant SMO674 (which includes an N-terminal hemagglutinin signal sequence and FLAG tag) along with GFP-tagged PKACα, we cloned a SMO674-IRES-PKACαGFP cassette into the pCDNA5-FRT-TO vector. Untagged human and mouse PKACα constructs in the vector pRSETb were previously described^85^. All mutant DNA sequences were prepared via PCR-based mutagenesis, cloned into their respective vector backbones via Gibson assembly, and verified by Sanger sequencing.

### Peptide synthesis

SMO PKI peptides (standard = SADVSSAWAQHVTKMVARRGAILP; extended = SADVSSAWAQHVTKMVARRGAILPQDVSVTPVATPVPP) were prepared via standard solid-phase synthesis by either GenScript or Elim Bio, purified via reversed-phase HPLC to ≥85% purity, and the sequence / molecular mass confirmed by mass spectrometry. For fluorescence polarization assays, a FAM fluorophore and 3xPEG linker were added to the N-terminus of the standard SMO PKI peptide during synthesis. PKIα(5-24) Peptide (TTYADFIASGRTGRRNAIHD) was synthesized as described for the SMO-PKI peptides by GeneCust (Boynes, France).

### Fluorescence polarization assays

Human or mouse PKA-Cα subunits were purified as previously described^87^. In addition, the protein obtained from the S200 was further purified via cation exchange chromatography. For this, the S200 peak fractions were pooled and dialyzed overnight into MonoS buffer (20 mM KH_2_PO_4_ pH 6.5, 5 mM DTT) before loading onto a MonoS cation exchange column. PKA-Cα was eluted with a gradient of 0-300mM KCl in MonoS buffer. Binding of a SMO peptide containing the PKI motif was investigated by fluorescence polarization (FP). Assay buffer for all FP experiments consisted of 50 mM MOPS pH 7, 35 mM NaCl, 10 mM MgCl_2_, 1 mM DTT and 0.005% Triton X-100 with the addition of 1 mM ATP in the PKA-C titrations. Experiments were performed by adding 50 μl of the titration component (PKA-C or ATP) to 150 μl of FAM-SMO containing solution. For SMO/PKA-C binding experiments, a two-fold dilution series from 16uM to 0uM of either mouse or human PKA-Cα was added to 40 nM FAM-SMO. To assess the ATP dependence of SMO binding to PKA-C, a titration with varying the ATP concentration from 0 to 1 mM (human PKA-Cα) or 12.8 μM (mouse PKA-Cα) was used with keeping PKA-C at 3 μM and FAM-SMO at 25 nM. Readings were taken with a Tecan Genios plate reader using black flat bottom 96-well Costar plates. Each experiment was carried out in at least triplicate with FAM readings at 485 nm excitation and 535 nm emission.

### Peptide arrays

Peptide arrays were synthesized with a MultiPep Flexible Parallel Peptide Synthesizer (Intavis Bioanalytical Instruments, Germany). The blots were probed with mouse or human PKA Cα protein, purified as described above for “Fluorescence polarization assays”. Unless noted otherwise all steps were carried out at room temperature. The blots were initially soaked with 100% ethanol for 5 minutes followed by 5 times 5-minute washes with water. All subsequent washes were carried out for 5 minutes 5 times with TTBS. The membranes were washed with TTBS, blocked with 5% milk in TTBS for 1 hour and washed again with TTBS. They were then soaked with TTBS containing 2 μg PKA-Cα, 5% milk, 10 mM MgCl_2_ and 1mM ATP overnight at 4° C. The next day the blots were washed, blocked with 5% milk in TTBS and washed again before incubation overnight at 4° C with TTBS containing 5% milk and a primary PKA-C antibody (generated in-house previously). The next day the blots were washed, then incubated with an HRP-conjugated anti-rabbit antibody (Prometheus™) in 5% milk TTBS for one hour followed by a final round of washes with TTBS. The membranes were finally covered with SuperSignal West Pico PLUS Chemiluminescent Substrate for detection of HRP (Thermo Scientific # 34580) and imaged with a ChemiDoc MP Imaging System from BIO RAD. A representative array image (representative of two separate trials) is shown; we excluded from analysis any spots that were not observed consistently across replicates.

### Surface plasmon resonance (SPR)

Human PKA-Cα was overexpressed in *E. coli* BL21(DE3) cells after induction with 0.4 mM IPTG for 16 h at RT using the expression vector pRSETb-hPKACα and then purified by affinity chromatography using an IP20-resin as described earlier^88^. GST-PKI, GST-SMO pCT wt and WRR mutant were overexpressed in *E. coli* BL21(DE3) for 16 hr at room temperature, and the GST fusion proteins were purified using Protino gluthathione agarose 4B according to the manufacturer’s instruction (Macherey-Nagel). SPR interaction studies were performed according to previous studies^46,85^. Briefly, the interaction studies were performed in running buffer (20 mM MOPS, pH 7.0, 150 mM NaCl, 50 μM EDTA, 0.005% P20 surfactant) at 25 °C using a Biacore 3000 instrument (GE Healthcare). For measurements involving ATP/MgCl_2_, the buffer was supplemented with 1 mM ATP and 10 mM MgCl_2_. Polyclonal anti-GST antibody (Carl Roth, 3998.1) was covalently immobilized to all four flow cells of a CM5 sensorchip (GE Healthcare) to a level of 8,000 response units (RU) via standard NHS/EDC amine coupling. Each measurement cycle started with the sequential capture of 60-130 RU of GST-PKI and 500-1,000 RU of GST-SMO-RLG wt and GST-SMO-RLG WRR mutant on separate flow cell (flow rate 10 µL/min). Interaction analysis was then initiated by the injection of increasing concentrations of human PKA Cα (156 nM – 10 µM) at a flow rate of 30 μL/min for 120 or 240 s (association) followed by 120 or 240 s dissociation with buffer without PKA Cα. Nonspecific binding was removed by subtracting SPR signals from a blank flow cell (without GST-protein) and additional buffer blank runs without PKA Cα (double referencing) employing BIAevaluation Software 4.1.1 (Cytiva, Marlborough, MA, USA). After each cycle, the sensorchip was regenerated by injecting up to 5 times with 10 mM glycine, pH 1.9 or 2.2, to remove the GST-fusion proteins from the antibody surfaces until the baseline level was reached. Steady-state analysis was performed with GraphPadPrism with a one-site binding (hyperbola) model as previously described^85^.

### NMR spectroscopy

Recombinant human PKA-Cα (PRKACA, uniprot P17612) was expressed and purified as reported^89,90^. Briefly, transformed *E*.*coli* BL21 (DE3) pLysS (Agilent) cells were cultured in deuterated (^2^H) M9 minimal medium supplemented with ^15^NH_4_Cl. Protein overexpression was initiated by adding 0.4 mM of isopropyl β-D-1-thiogalactopyranoside (IPTG) and carried out for 12 hours at 20 °C. The collected cell pellet was then resuspended in 50 mM Tris-HCl pH 8.0, 30 mM KH_2_PO_4_, 200 mM NaCl, 5 mM β-mercaptoethanol, 0.15 mg/ mL lysozyme, 200 μM ATP, DnaseI, and protease inhibitor (Sigma) and pass through a French press (2 times). The cell resuspension was then cleared by centrifugation (18.000 rpm for 45 minutes), and the supernatant was incubated overnight with Ni^2+^-NTA resin (Thermo Fisher). His-tagged PKA-Cα was eluted using 50 mM Tris-HCl pH 8.0, 30 mM KH_2_PO_4_, 100 mM NaCl, 5 mM β-mercaptoethanol, 1 mM phenylmethylsulfonyl fluoride (PMSF) supplemented with 200 mM of imidazole. The His-tag was removed during an overnight dialysis step performed in 20 mM KH_2_PO_4_ (pH 6.5), 25 mM KCl, 5 mM β-mercaptoethanol, 0.1 mM PMSF, using stoichiometric quantities of recombinant tobacco etch (TEV) protease. Finally, cationic exchange chromatography was performed to separate the three different isoforms of PKA-Cα, representing the three different phosphorylation states of the kinase, using a linear gradient of KCl in 20 mM KH_2_PO_4_ at a pH of 6.5 (HiTrap Q-SP column, GE Healthcare Life Science). All the NMR experiments were performed using PKA-Cα isoform II, corresponding to phosphorylation at S10, T197, and S338 residues^91^. The purity of the protein preparation was tested using SDS-PAGE electrophoresis and Mass spectrometry analysis (purity >97%).

A portion of the lyophilized extended SMO PKI peptide (see above) was resuspended in 20 mM KH_2_PO_4_ (pH 6.5), 90 mM KCl, 10 mM MgCl_2_, 10 mM DTT, 1 mM NaN_3_, to a final concentration of 4 mM. The NMR experiments were performed on 100 μM uniformly ^2^H, ^15^N-labeled PKA-Cα sample in 20 mM KH_2_PO_4_, 90 mM KCl, 10 mM MgCl_2_, 10 mM DTT, 1 mM NaN_3_ at pH of 6.5 and saturated with 12 mM of a non-hydrolyzable ATP analog (ATPγN). Modified [^1^H-^15^N]-TROSY-HSQC spectra were acquired on a Bruker Advance III spectrometer operating at a proton frequency of 850 MHz, equipped with a TCI cryoprobe, at an acquisition temperature of 300K. First, a spectrum of ATPγN-saturated PKA-Cα complex (PKA-Cα/ATPγN) was recorded with 2048 (proton) and 128 (nitrogen) complex points. Stoichiometric amounts of SMO PKI peptide (hereafter referred to as SMO-PKI) were then added to the PKA-Cα/ATPγN complex until saturation (1:0, 1:1, 1:2, and 1:4 SMO-PKI:PKA-C/ATPγN molar ratio), with a concentration of SMO-PKI ranging from 0.1 to 0.4 mM. All the spectra were processed using NMRPipe^92^, and visualized using NMRFAM-SPARKY software^93^. Combined chemical shift perturbation (CSPs) were calculated using the ^1^H and the ^15^N chemical shift derived from the PKA-Cα/ATPγN complex and the 1:4 SMO-PKI:PKA-C/ATPγN complex. The CSPs was calculated using the following equation^94^:

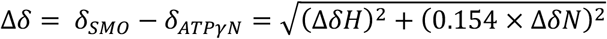

The PKIα(5-24) CSPs were originally published earlier^90^ and are included here for reference.

### Purification of SMO pCT for PKA-C activity assays

MBP-His_8_-SUMO-tagged SMO pCT (SMO pCT(565-657) / pHTSHP) was transformed into BL21 (DE3) *E coli* and grown to OD_600_= ∼0.5-1.0 in 1 L terrific broth (TB) with ampicillin at 37°C in 2800ml baffled Fernbach flasks until. IPTG was then added to 0.4mM and the temperature lowered to 18°C for 18 hours. Cells were harvested by centrifugation, snap-frozen in liquid nitrogen, and stored at −80°C. Cell pellets were resuspended in 20ml binding buffer (50mM Tris pH 8.0, 300mM NaCl, 10mM imidazole, protease inhibitors (Pierce)) at 4°C with stirring. Lysozyme was added at 1mg/ml as well as benzonase (10,000X, Sigma), and samples were lysed by sonication. Lysates were clarified by centrifugation at 40,000 x g, 30 min, 4°C. Supernatant was run by gravity over a column of 6ml NiNTA affinity resin (Qiagen) pre-equilibrated in binding buffer. The column was washed twice with 5 column volumes of wash buffer (50mM Tris pH 8.0, 300mM NaCl, 25mM imidazole) and eluted with four successive 1 column volumes of elution buffer (50mM Tris pH 8.0, 300mM NaCl, 250mM imidazole). Protein content of fractions was estimated by Quickstart Bradford assay and purity determined by SDS-PAGE. Protein-rich fractions were pooled and cleaved by the addition of 1.4mg Ulp1 enzyme (prepared in-house) and 1mM DTT for 1 hour at room temperature. After incubation, the samples were dialyzed overnight against dialysis buffer (50mM HEPES pH 8.0, 150mM NaCl, 7mM β-mercapthoethanol. The completed digestion reaction was applied to NiNTA resin, washed twice with 3 column volumes of dialysis buffer, and eluted with 3 column volumes of elution buffer. The flow-through and wash fractions were pooled, centrifugated at 20,000 x g, 5 min, 4°C to pellet any insoluble material, and futher purified by loading onto a 1ml HiTrap HP SP cation exchange column pre-equilibrated in low-salt buffer (50mM HEPES pH8.0, 150mM NaCl). Column was washed extensively with low salt buffer and protein was eluted with a gradient to high-salt buffer (50mM HEPES pH 8.0, 1.2M NaCl). Fractions containing the cleaved SMO pCT were pooled, concentrated, and injected onto a Superdex 200 (10/300) gel filtration column equilibrated in gel filtration buffer (50 mM HEPES pH 7.5, 300 mM NaCl). The peak fractions were collected and analyzed by SDS-PAGE. Fractions containing intact pCT were pooled, snap-frozen in liquid nitrogen, and stored at −80°C. For the experiments in Fig. 2d, wild-type or mutant pCT proteins were purified by NiNTA affinity as described above and used directly in PKA-C activity assays.

### *In vitro* PKA-C activity assays

For *in vitro* PKA-C activity assays the recombinant mouse PKA catalytic subunit (Cα) was overexpressed in *E. coli* BL21(DE3) cells after induction with 0.4 mM IPTG for 14 hr at room temperature using the expression vector pRSETb / mPKACα and then purified by affinity chromatography using an IP20-resin as described earlier^88^. PKA catalytic activity was assayed using a coupled spectrophotometric assay as described previously^42^. Briefly, the reaction mixture contained 100 mM MOPS (pH 7), 10 mM MgCl_2_, 100 µM ATP, 1 mM phosphoenolpyruvate, 15 U/mL lactate dehydrogenase, 70 U/mL pyruvate kinase, 200 mM reduced nicotinamide adenine dinucleotide, 5 mM β-mercaptoethanol with 15-30 nM Cα and 260 µM Kemptide (LRRASLG; GeneCust) as a substrate. Formation of PKA-C:SMO complex (concentration of SMO peptide or recombinant SMO pCT protein are indicated in the figures) was carried out for 3 minutes at room temperature in the assay mixture. The apparent IC_50_ for SMO pCT were determined by fitting the concentration-dependent activity to a sigmoid dose-response model. All data were plotted as means of at least two independent experiments measured in duplicate each with standard deviation (SD).

### MALS

Purified SMO pCT was concentrated to 5 mg/ml and analyzed via SEC-MALS using a Superdex 75 gel chromatography column (GE Healthcare) equilibrated in gel filtration buffer (50 mM HEPES pH 7.5, 300 mM NaCl) with an in-line DAWN MALS detector (Wyatt Technology.)

### BRET

BRET in HEK293 or IMCD3 cells was performed as previously described^24^. Briefly, HEK293 or IMCD3 cells were transiently transfected with nanoluc-tagged SMO (0.3 μg) along with YFP-tagged βarrestin1, PKA-Cα, or PKA-RIα plasmids. Typically 0.1 μg of each BRET acceptor plasmid was used, with the following two exceptions: (1) Fig. 4b which examined a range of PKA-Cα DNA amounts as indicated in the figure legend; (2) Fig 3d which used 0.3 μg of PKA-Cα-YFP (corresponding to a 0.33-2.68% increase in the size of the endogenous PKA-C pool, as determined by quantitative immunoblotting^24^) or PKA-RIα-YFP (which displays no BRET with SMO even though it expresses at substantially higher levels than does PKA-Cα-YFP^24^); Cells were replated in poly-D-lysine coated white opaque 96-well plates, loaded with 5 μM coelenterazine h (HEK293) or 10 μM furimazine (IMCD3), and analyzed for BRET on a Tecan Spark multimode plate reader. The background signal from cells expressing nanoluc-tagged SMO without BRET acceptor was subtracted from all measurements. For all BRET assays, data represent mean ± SEM from triplicate wells, and data are representative of at least two independent experiments.

### Flow cytometry

Cell surface expression of wild-type and mutant SMO constructs was analyzed via flow cytometry as previously described^24^. Briefly, HEK293-Freestyle cells were infected with BacMam viruses encoding the indicated wild-type or mutant SMO674 constructs. 1-2 days later, live cells were stained with Alexa 647-conjugated M1 anti-FLAG antibody (1:1000, 5 minutes) and analyzed on a Cytoflex flow cytometer (Beckman Coulter).

### CREB reporter assays

CREB transcriptional reporter assays were performed as previously described^24^. Briefly, HEK293FT cells were transfected in 24-well plates with a 30:1 mixture of CRE-Firefly reporter (pGL4.29[luc2p/CRE/Hygro]) and constitutively expressing SV40-Renilla plasmids (20%(w/w)), PKA-Cα (0.625%(w/w)), along with the indicated wild-type or mutant SMO674 plasmids (24%(w/w)). A GFP expression plasmid was used to bring the total amount of DNA in each well to 250 ng. Two days later, reporter activity was measured via dual luciferase assay. Reporter activity is expressed as a ratio of Firefly/Renilla (relative luciferase units (RLU)) For all CREB assays, data represent mean ± SEM from triplicate wells, and data are representative of at least two independent experiments.

### Coimmunoprecipitation assay

SMO / PKA-C coimmunoprecipitation was performed as previously described^24^, with minor modifications. Briefly, 3 ml HEK293-Freestyle cells were infected with BacMam viruses encoding PKA-Cα-YFP the indicated wild-type or mutant SMO674 constructs and solubilized in low-salt solubilization buffer (20 mM HEPES pH 7.5, 150 mM NaCl, 0.1 mM TCEP, 0.5% GDN, 1 mM CaCl2•6H_2_O, protease inhibitor tablet) to prepare a whole-cell lysate. A portion of the lysate was reserved for SDS-PAGE analysis, and the remainder was incubated with FLAG affinity resin (10 μl settled resin per condition). After a one-hour incubation, resin was washed three times in low-salt wash buffer (20 mM HEPES pH 7.5, 150 mM NaCl, 0.1 mM TCEP, 0.05% GDN, 1 mM CaCl_2_•6H2O), and protein eluted in 40 μl of the same buffer supplemented with 5 mM EDTA and 0.2 mg/ml FLAG peptide. Proteins were separated by SDS-PAGE on a 4-20% Stain-Free TGX gel (BioRad), and total protein in lysate or eluate was visualized via Stain Free imaging. To detect PKA-Cα-YFP, inputs and eluates were transferred to PVDF membranes and processed for immunoblotting with anti-GFP antibodies as previously described^24^. In-gel Pro-Q Diamond assay to detect phosphoprotein was performed according to the manufacturer’s instructions, as previously described^24^.

### HEK293 imaging

Imaging and quantification of SMO / PKA-C colocalization in HEK293 cells were performed as previously described^24^, with minor modifications. Briefly, HEK293 Flp-in T-rex stable cell lines coexpressing GFP-tagged PKA-Cα with either wild-type or mutant FLAG-tagged SMO674 were plated onto μ−slide 8-well glass chamberslides (ibidi), and treated overnight with 1 μg/ml doxycycline (to induce SMO expression) and 1 μM SAG21k (to induce SMO activation). Live cells were subsequently stained for 5 min with an Alexa647-conjugated M1 anti-FLAG antibody (1:2000) followed by washing in HBSS, mounting, and visualization. Images were collected on a Leica SP8 laser scanning confocal microscope, using a 40x water immersion lens. All images were acquired with identical zoom / exposure / gain settings and processed identically in Fiji. SMO / PKA-C colocalization was determined by measuring the background-subtracted PKA-C (green) fluorescence at the membrane (GFP_membrane_ – GFP_cytoplasm_) and the background-subtracted SMO (red) fluorescence at the membrane (Alexa647_membrane_ – Alexa647_cytoplasm_), using four independent line-scans across the membrane in Fiji. The background-subtracted green fluorescence divided by the background-subtracted red fluorescence is referred to as “SMO / PKA-C colocalization” and reported in arbitrary units (AU).

### IMCD3 imaging

Imaging and quantification of SMO ciliary accumulation and SMO / PKA-C colocalization in IMCD3 cells were performed as previously described^24^. Briefly, to assess ciliary accumulation of wild-type or mutant SMO, IMCD3 cells were transiently transfected on coverslips with the indicated myc-tagged wild-type or mutant SMO / pGEN constructs, grown to confluency, fixed, and permeabilized. Coverslips were then stained with anti-myc (SMO) and anti-Arl13b (cilia) antibodies, along with DAPI to mark the nucleus. For quantitative assessment of SMO / PKA-C colocalization in live IMCD3 cilia, cell lines coexpressing mNeonGreen-tagged PKA-Cα with either wild-type or mutant FLAG-tagged SMO were plated onto μ−slide 8-well glass chamberslides (ibidi), grown to confluency, and treated overnight in low-serum medium with 1 μM SAG21k (to induce SMO activation and ciliary accumulation.) A previously described cell line coexpressing mNeonGreen-tagged Nbβ2AR80 with wild-type FLAG-tagged SMO^24^ was used as a negative control. Live cells were subsequently stained for 5-10 min with an Alexa647-conjugated M1 anti-FLAG antibody (1:1000) and Hoechst counterstain, followed by washing in HBSS, mounting, and visualization. Cells were imaged on a Leica SP8 laser scanning confocal using a 40x water immersion lens. Three-dimensional reconstructions of Z-stacks were performed in Fiji using the 3D Viewer plugin. Quantification of SMO (red) and PKA-C/Nbβ2AR80 (green) staining was performed using CiliaQ^95^ and reported as a ratio of the green fluorescence in each cilium, normalized to the red fluorescence (“Colocalization with SMO in cilia (AU)”), as previously described^24^. All images were acquired with identical zoom / exposure / gain settings.

### GLI reporter assays

GLI transcriptional reporter assays were performed as previously described^24^. Briefly, *Smo*^-/-^ MEFs were transfected with a 30:1 mixture of 8xGli-Firefly and SV40-Renilla plasmids (50% (w/w)), along with the indicated full-length wild-type or mutant SMO constructs (in a pGEN vector, with C-terminal myc tag) (2%(w/w)), and GFP to adjust the amount of DNA to 250 ng/well. Cells were cultured to confluency, shifted to 0.5% FBS-containing medium, and treated with control or ShhN conditioned medium (1:20 dilution) for 2 days. Reporter activity was then measured via dual luciferase assay. For all GLI assays, data represent mean ± SEM from triplicate wells, and data are representative of at least two independent experiments.

### Zebrafish embryological studies

Zebrafish mRNA injection, fixing, mounting, and staining of muscle cell markers were all performed as previously described^24^, except that a far-red secondary antibody (donkey anti-rabbit 649, Jackson Laboratory, 1:500) was used instead of a red secondary antibody during immunohistochemistry.

### Software

PDB files were viewed using Pymol. Graphs, curve fitting, and statistical analysis was performed in GraphPad Prism. Alignments were generated using CLUSTAL Omega. Graphics were generated using BioRender or Adobe Illustrator.

## ACKNOWLEDGMENTS

We thank J. Zalatan for making us aware of the parallels between SMO / PKA-C regulation in the Hh pathway and LRP / GSK-3β regulation in the Wnt pathway. We thank S. Lusk and K. Kwan for providing *smo-*null zebrafish (*smo*^hi1640^), and D. Klatt Shaw and D. Grunwald for sharing advice and reagents regarding zebrafish immunohistochemistry. We thank J. Müller and S. Kasten for excellent technical assistance. We thank the Johnson Foundation Structural Biology and Biophysics Core at the Perelman School of Medicine (Philadelphia, PA) for performing SEC-MALS analyses. We thank D. Julius, K. Basham, A. Manglik, M. He, and S. Nakielny for providing feedback on this manuscript. B.R.M. acknowledges support from the 5 for the Fight Foundation and a Cancer Center Support Grant Pilot Project Fund from the Huntsman Cancer Institute. This work was supported by the funding line Future (PhosMOrg) of University of Kassel (F.W.H.), and NIH grants R01GM100310-08 (G.V.), 1R35GM130389 (S.S.T.), 1R03TR002947 (S.S.T.) and 1R35GM133672 (B.R.M.).

## AUTHOR CONTRIBUTIONS

J.T.H. designed, executed, and interpreted CREB and GLI reporter assays. C.D.A. designed, executed, and interpreted HEK293 BRET assays. J.B. designed, executed, and interpreted fluorescence polarization studies and peptide array studies. D.B. designed, executed, and interpreted *in vitro* PKA-C activity assays and SPR studies (the latter with assistance from J.W.B.). I.B.N. developed SMO pCT purification approaches and purified this domain for *in vitro* PKA-C activity assays. C.O. designed, executed, and interpreted NMR studies. D.S.H. designed, executed, and interpreted zebrafish embryology studies. J-F.Z. designed, executed, and interpreted co-immunoprecipitation and IMCD3 BRET assays. J.L.C. designed, executed, and interpreted HEK293 confocal imaging studies. L.V. performed initial fluorescence polarization studies. C.C.K. collaborated with J.B. to develop SMO peptide arrays. V.L.R-P. provided advice and guidance on mutagenesis experiments to disrupt SMO / PKA-C interactions. S.S.T. and B.R.M. conceived the project. G.V., F.W.H., S.S.T., and B.R.M. interpreted data and provided overall project supervision. B.R.M. performed IMCD3 ciliary imaging studies and wrote the manuscript with assistance from J.T.H., C.D.A., and I.B.N.

## COMPETING FINANCIAL INTERESTS

The authors declare no competing financial interests.

### Correspondence and requests for materials

should be addressed to S.S.T. or B.R.M.

## FIGURES

**Extended Data Fig.1:**
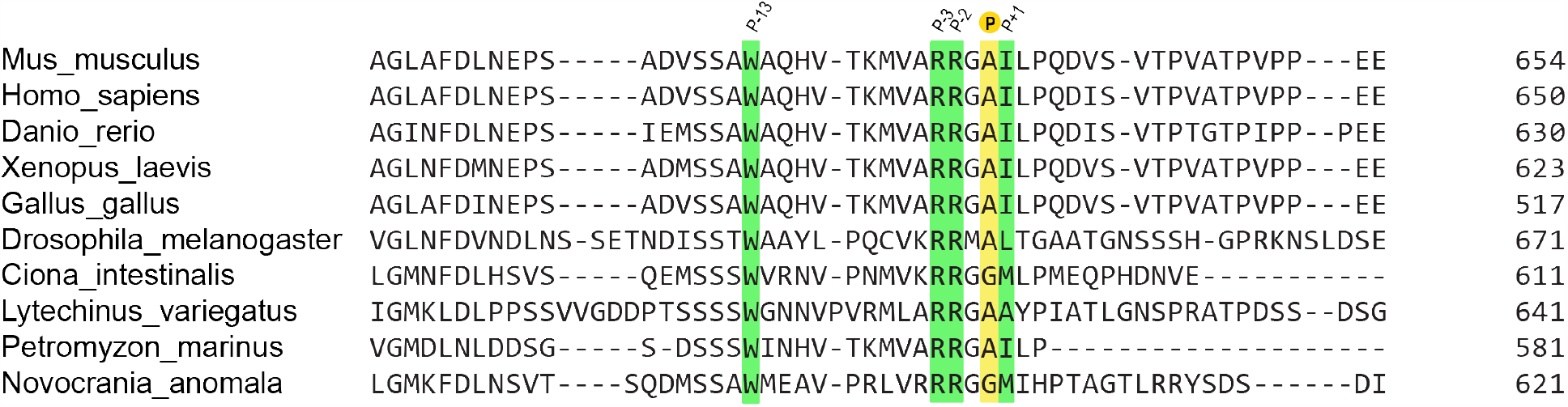
Sequence alignment of SMO PKI motif. Extended alignment of a portion of the pCT from the indicated SMO orthologs, with key PKI motif residues colored as in Fig. 1a.

**Extended Data Fig. 2:**
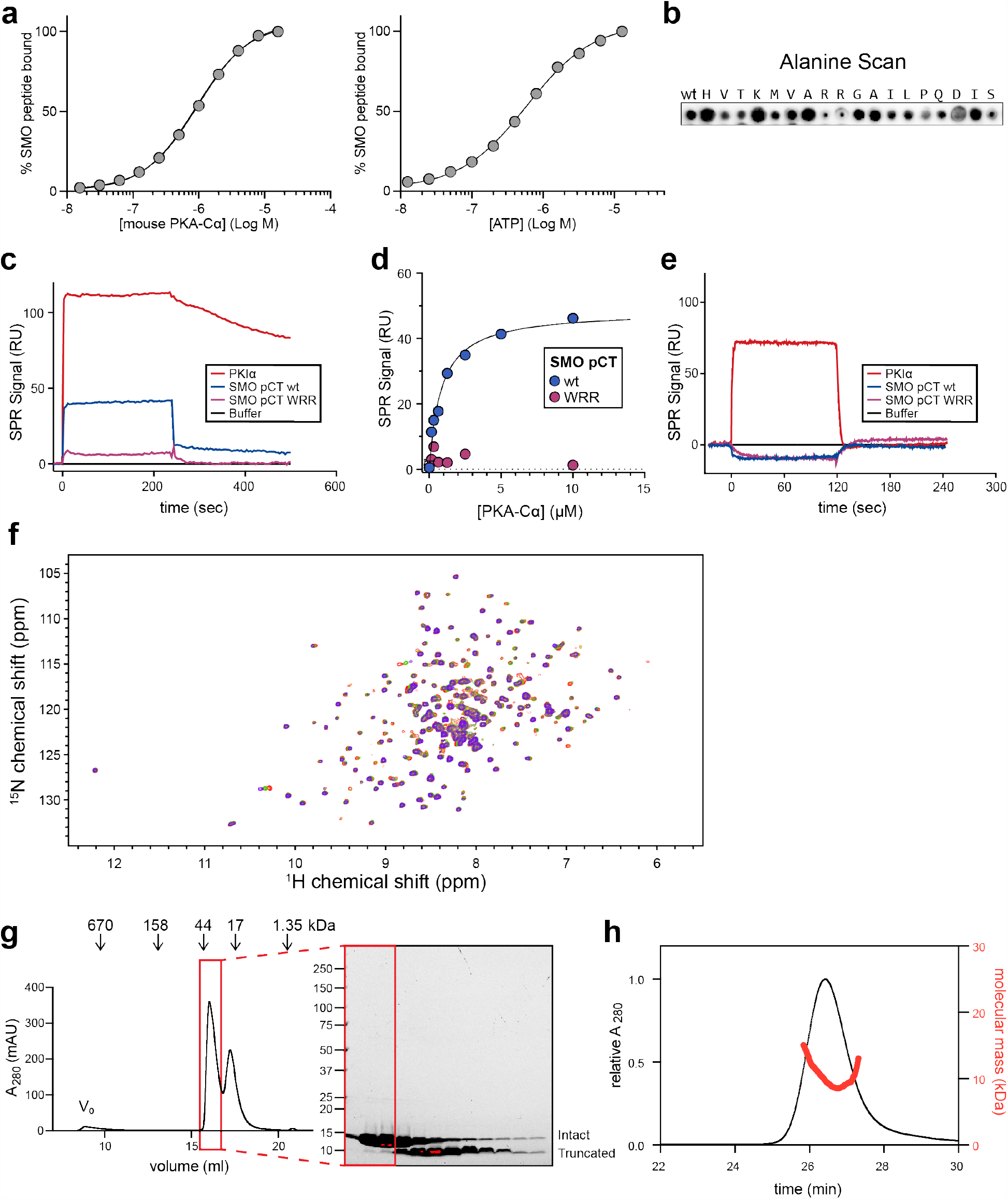
Further binding and peptide array studies, SPR sensorgrams, NMR spectra, and SMO pCT purification strategy. **a**, Fluorescence polarization assays using mouse PKA-Cɑ, performed as in Fig. 1b. **b**, Peptide array, performed as in Fig. 1c, but with individual residues in the human SMO PKI motif mutated to alanine. **c**, SPR sensorgram for 5 μM PKA-Cɑ binding to GST-tagged wild-type (blue) or WRR mutant (purple) SMO pCT, or a PKIɑ positive control (red), in the presence of ATP and MgCl_2_. **d**, Steady-state analysis of binding interactions between human PKA-Cɑ and a recombinant wild-type (wt, blue) or mutant (WRR, magenta) SMO pCT, as assessed by SPR. **e**, SPR sensorgram, performed as in **c**, but with ATP and MgCl_2_ omitted from the buffer. PKA-Cɑ was present at 2.5 µM. Note that although removal of ATP and MgCl_2_ does not completely eliminate steady-state binding to the PKIɑ positive control, it dramatically accelerates the dissociation rate, as expected. **f**, [^1^H, ^15^N] heteronuclear single quantum coherence (HSQC) spectra used to calculate CSP values. Spectra were acquired from PKA-Cɑ / ATPɑN complexed with a SMO peptide at 1:0 (red), 1:1 (yellow), 1:2 (green), or 1:4 (purple) molar ratios. CSP values were calculated from the spectrum corresponding to a 1:4 molar ratio and plotted in Fig. 2a. **g**, Purification of SMO pCT domain from *E. coli*. Following size exclusion chromatography (left), the purified protein was analyzed by SDS-PAGE (right). The SMO pCT elutes as two peaks, the earlier of which is the intact pCT and the later of which corresponds to a truncated fragment (data not shown). Fractions containing the intact pCT (red box) were pooled and used for subsequent experiments. **h**, Multi-angle light scattering coupled with size exclusion chromatography (SEC-MALS) was used to determine the protein oligomeric state for the pooled fractions indicated in **g**. The average molecular mass was calculated as M_w_ = 11.25 +/-2.1 kDa, close to the predicted molecular mass for a monomer (10.1 kDa).

**Extended Data Fig. 3:**
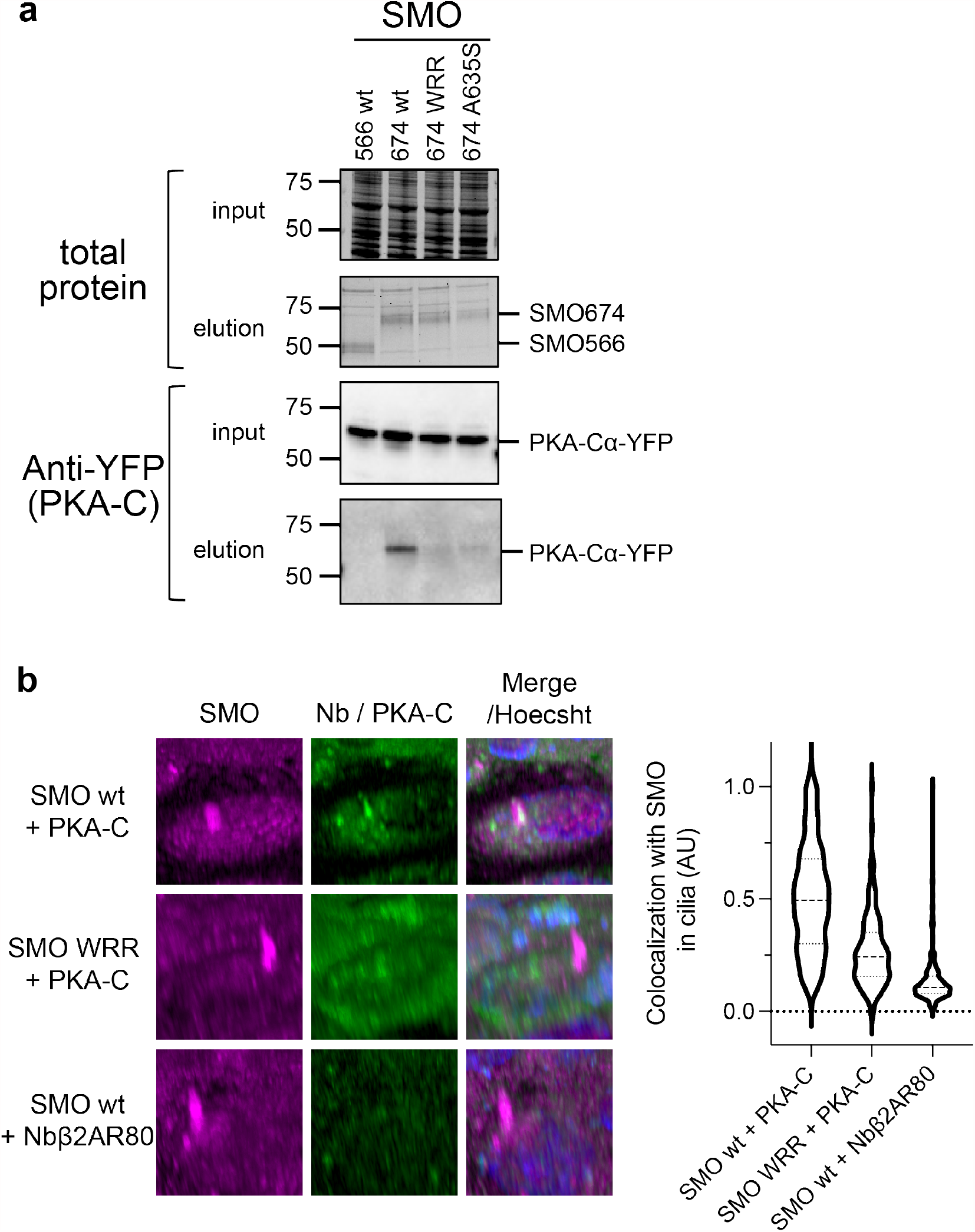
Coimmunoprecipitation studies and ciliary colocalization studies to assess SMO / PKA-C interactions. **a**, Coimmunoprecipitation of PKA-Cɑ-YFP with the indicated FLAG-tagged wild-type or mutant SMO constructs was assessed using FLAG chromatography from lysates of transfected HEK293 cells. **b**, Left, Colocalization of FLAG-tagged wild-type or mutant SMO674 (magenta) with mNeonGreen-tagged PKA-Cɑ (green) in ciliated IMCD3 cells stably expressing both constructs and treated with a SMO agonist, SAG21k, to induce SMO ciliary localization. Cilia are marked by the SMO stain. mNeonGreen-tagged Nbβ2AR80 (which does not bind SMO^24^) serves as a negative control. 3D reconstructions from Z-stacks of confocal live-cell images are shown. Right, quantification of microscopy studies (n=142-244 cilia per condition). See Extended Data Table 1 for statistical analysis.

**Extended Data Fig. 4:**
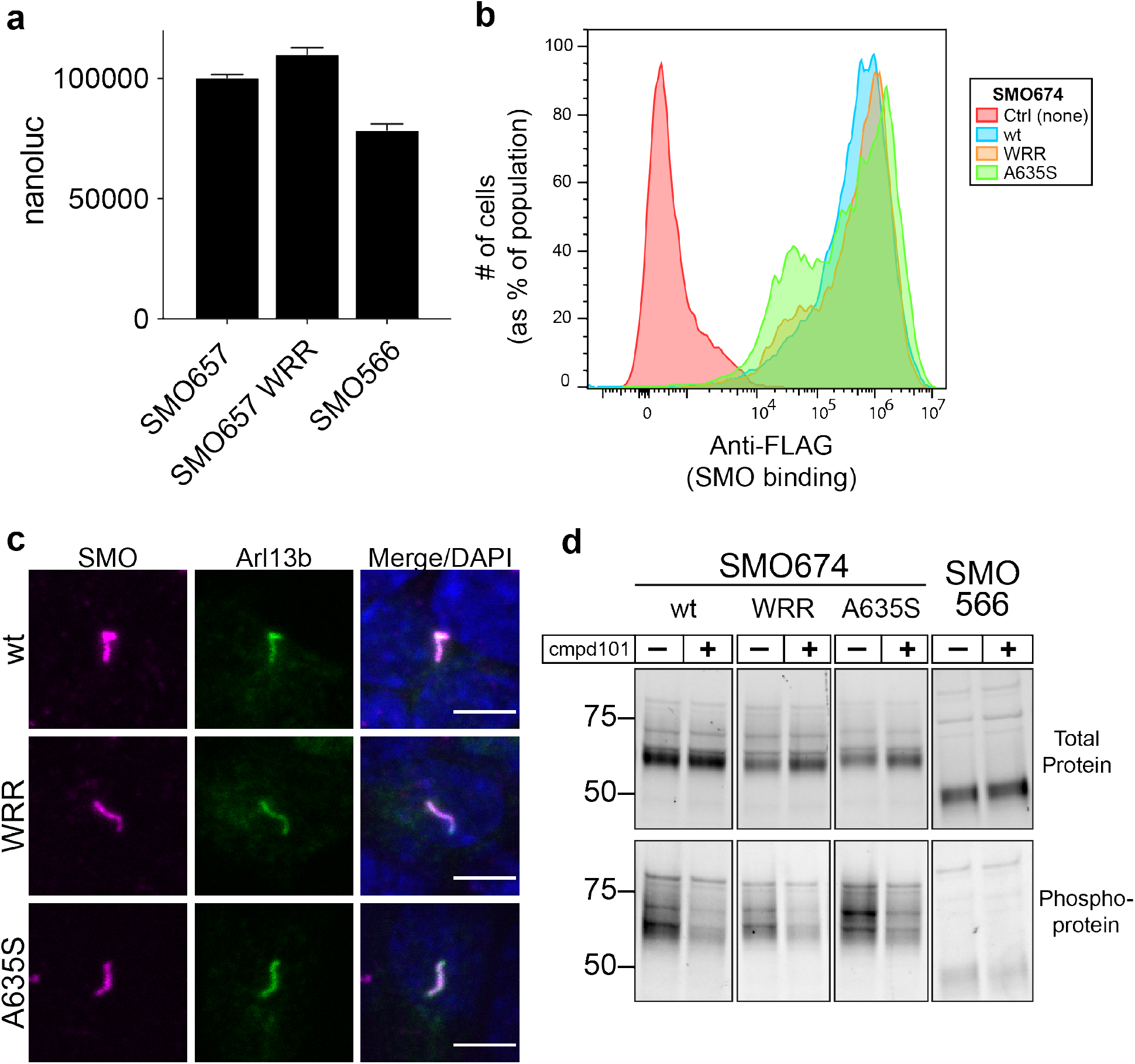
Controls for SMO / PKA-C binding, colocalization, and signaling studies. **a**, Expression levels of SMO constructs in Fig. 3a, assessed by whole-cell nanoluc measurements. **b**, Surface levels of N-terminally FLAG-tagged wild-type or mutant SMO674 constructs were quantified via expression in HEK293 cells followed by FLAG staining and flow cytometry. Mock-infected cells stained with FLAG antibody (red) serve as a negative control. **c**, Ciliary localization in IMCD3 cells of myc-tagged wild-type or mutant SMO proteins (magenta). Cilia were visualized with Arl13b antibody (green). Scale bar = 5 µm. **d**, GRK2/3-dependent phosphorylation of FLAG-tagged wild-type or mutant SMO674 constructs was determined via expression in HEK293 cells treated with or without the GRK2/3 inhibitor cmpd101, followed by FLAG purification. Levels of total and phosphorylated SMO were assessed by Stain Free imaging and ProQ Diamond fluorescence, respectively. SMO566, which is not phosphorylated by GRK2/3 (as it does not contain the C-tail and therefore lacks all previously mapped physiological GRK2/3 phosphorylation sites^24^), serves as a negative control.

**Extended Data Fig. 5:**
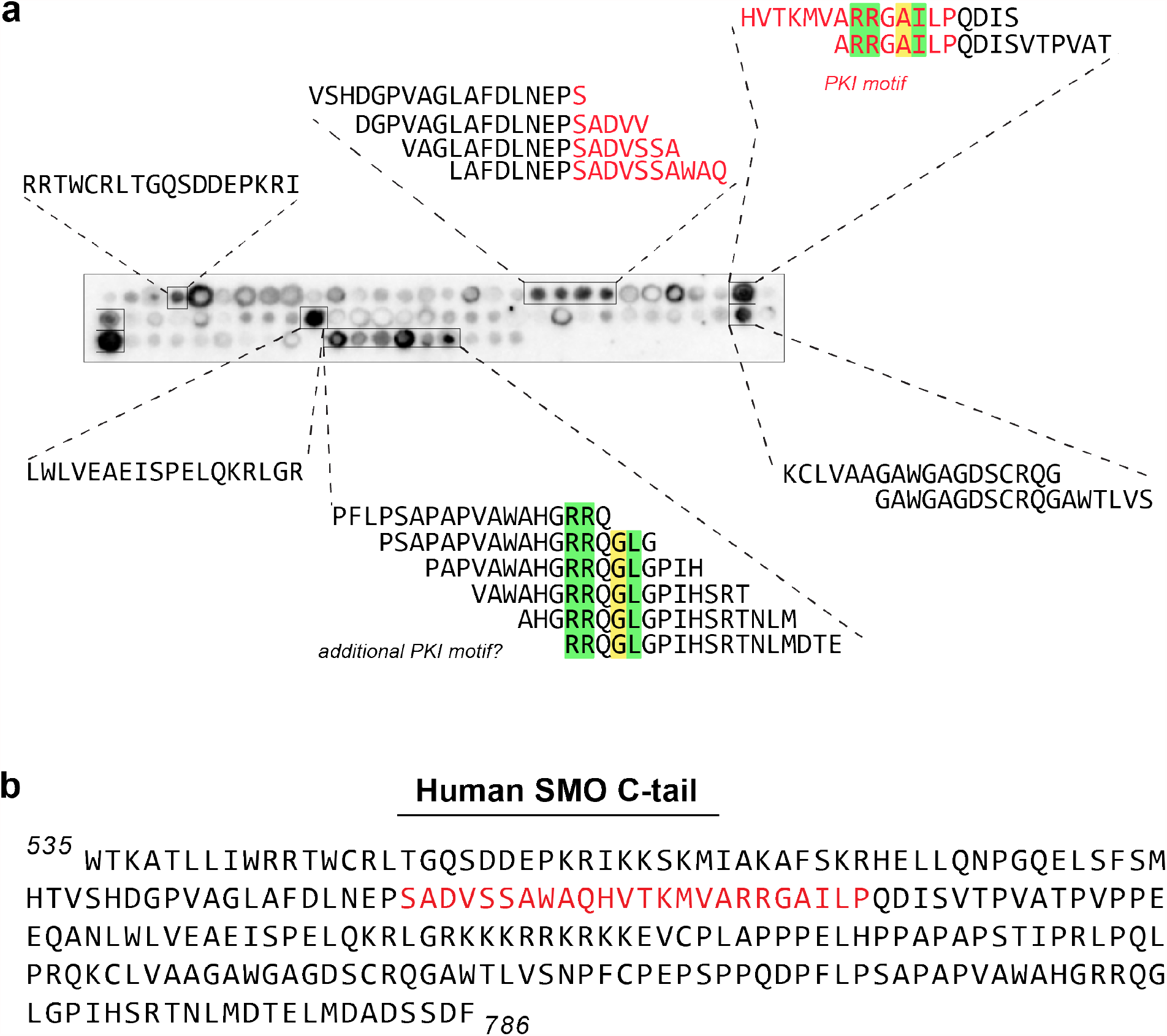
Complete data set from SMO C-tail peptide array studies. **a**, The same SMO tiled peptide array from Fig. 4c, but including the sequences of all positive hits in each array cluster. **b**, Complete human SMO C-tail sequence used to create the peptide array. In **a**,**b**, the SMO PKI motif identified in the pCT is indicated in red. Key residues in this PKI motif, along with ones in the candidate PKI motif in the dCT, are colored as in Fig. 1a.

**Extended Data Fig. 6:**
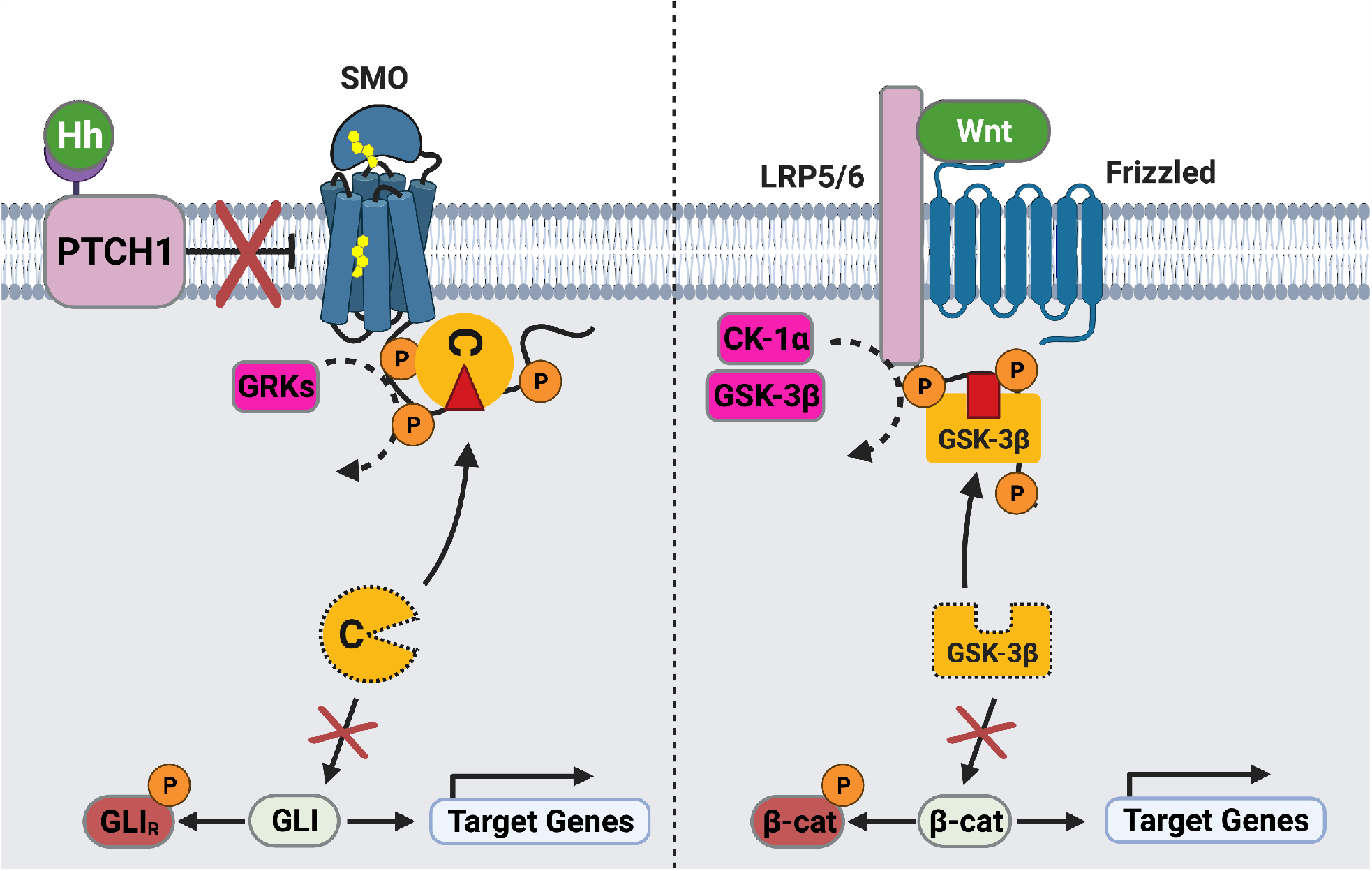
Similarity between signal transduction mechanisms in the Hh and Wnt pathways. Schematic diagram of transmembrane signal transduction in the Hh (left) and Wnt (right) pathways. During Hh signal transduction, active SMO is phosphorylated on its cytoplasmic tail by GRK2/3, triggering membrane sequestration and inhibition of PKA-C, and ultimately stabilization and activation of GLI. During Wnt signal transduction, active LRP5/6 is phosphorylated on its cytoplasmic tail by glycogen synthase kinase (GSK)-3β and casein kinase (CK)-1ɑ, triggering membrane sequestration and inhibition of GSK-3β, and ultimately stabilization and activation of β-catenin^74-77^. Note that this is a simplified and highly schematized diagram and is not intended to be comprehensive; many other components of both pathways (for example, the destruction complex in which GSK-3β and β-catenin reside) are omitted in order to highlight mechanistic similarities between the underlying transmembrane signaling mechanisms.

**Extended Data Table 1.** Tests of statistical significance for all figures.

| Figure | Condition 1 | Condition 2 | Significant? | p-value |
| --- | --- | --- | --- | --- |
| 2b | multiple comparison | multiple comparison | yes | <0.001<sup>a</sup> |
| 2d | None | wt | yes | <0.0001<sup>b</sup> |
| 2d | WRR | wt | yes | <0.0001<sup>b</sup> |
| 2d | None | WRR | no | n.s.<sup>b</sup> |
| 3a | SMO657 wt: βarrestin1 | SMO657 wt: NbSmo2 | yes | 0.00053 |
| 3a | SMO657 wt: βarrestin1 | SMO657 wt: PKA-C | yes | 0.00438 |
| 3a | SMO657 wt: PKA-C | SMO657 WRR: PKA-C | yes | 0.001553 |
| 3a | SMO657 wt: PKA-C | SMO566 wt: PKA-C | yes | 0.003181 |
| 3a | SMO657 WRR: βarrestin1 | SMO657 WRR: PKA-C | no | 0.051139 |
| 3a | SMO657 WRR: βarrestin1 | SMO657 WRR: NbSmo2 | yes | 0.000165 |
| 3a | SMO566 wt: βarrestin1 | SMO566 wt: PKA-C | no | 0.22268 |
| 3a | SMO566 wt: βarrestin1 | SMO566 wt: NbSmo2 | yes | 0.00162 |
| 3b | SMO674: wt | SMO674: WRR | yes | <0.0001 |
| 3c | None | PKA-C | yes | 0.000723 |
| 3c | PKA-C | PKA-C + SMO674 wt | yes | 0.000995 |
| 3c | PKA-C | PKA-C + SMO674 WRR | no | 0.267104 |
| 3c | PKA-C | PKA-C + SMO674 A635S | no | 0.544481 |
| 3c | PKA-C + SMO674 wt | PKA-C + SMO674 WRR | yes | 0.004314 |
| 3c | PKA-C + SMO674 wt | PKA-C + SMO674 A635S | yes | 0.000513 |
| 3d | SMO wt: PKA-R | SMO wt: PKA-C | yes | 0.012831 |
| 3d | SMO wt: PKA-C | SMO WRR: PKA-C | yes | 0.01315 |
| 3d | SMO WRR: PKA-R | SMO WRR: PKA-C | no | 0.113776 |
| 3e | Ctrl.: Ctrl | Neg.: ShhN | no | 0.63116 |
| 3e | SMO wt: Ctrl | SMO wt: ShhN | yes | 0.010704 |
| 3e | SMO wt: ShhN | SMO WRR: ShhN | yes | 0.010346 |
| 3e | SMO wt: ShhN | SMO A635S: ShhN | yes | 0.010945 |
| 4b | βarrestin1: SMO WRR | PKA-C (0.04 μg): SMO WRR | no | 0.885586 |
| 4b | βarrestin1: SMO WRR | PKA-C (0.08 μg): SMO WRR | yes | 0.001529 |
| 4b | βarrestin1: SMO WRR | PKA-C (0.12 μg): SMO WRR | yes | 0.000368 |
| 4b | βarrestin1: SMO WRR | PKA-C (0.16 μg): SMO WRR | yes | 0.000109 |
| 4b | βarrestin1: SMO WRR | PKA-C (0.20 μg): SMO WRR | yes | 0.000085 |
| 4b | PKA-C (0.04 µg): SMO wt | PKA-C (0.04 µg): SMO WRR | yes | 0.004444 |
| 4b | PKA-C (0.08 µg): SMO wt | PKA-C (0.08 µg): SMO WRR | yes | 0.000502 |
| 4b | PKA-C (0.12 µg): SMO wt | PKA-C (0.12 µg): SMO WRR | yes | 0.001341 |
| 4b | PKA-C (0.16 µg): SMO wt | PKA-C (0.16 µg): SMO WRR | yes | 0.004344 |
| 4b | PKA-C (0.20 µg): SMO wt | PKA-C (0.20 µg): SMO WRR | yes | 0.000029 |
| 4b | PKA-C (0.04 µg): SMO566 | PKA-C (0.04 µg): SMO WRR | yes | 0.00368 |
| 4b | PKA-C (0.08 µg): SMO566 | PKA-C (0.08 µg): SMO WRR | no | 0.180776 |
| 4b | PKA-C (0.12 µg): SMO566 | PKA-C (0.12 µg): SMO WRR | yes | 0.014786 |
| 4b | PKA-C (0.16 µg): SMO566 | PKA-C (0.16 µg): SMO WRR | yes | 0.006063 |
| 4b | PKA-C (0.20 µg): SMO566 | PKA-C (0.20 µg): SMO WRR | yes | 0.001566 |
| ED 3b | SMO wt + PKA-C | SMO WRR + PKA-C | yes | <0.0001 |
| ED 4a | SMO657 | SMO657 WRR | no | 0.057394 |

<sup>a</sup> To check for significant differences between PKA-C lacking peptide (control) and each peptide concentration a one-way ANOVA with Tukey's multiple comparison was performed. All comparisons show $p \leq 0.0001$ (\*\*\*\*) significance except the comparison between 125µM standard vs extended peptide ( $p \leq 0.001$ , \*\*\*).
<sup>b</sup> To check for significant differences between PKA lacking peptide (control) and the indicated recombinant pCT fragment a one-way ANOVA with Tukey's multiple comparison was performed (\*\*\*\* $\triangleq p \leq 0.0001$ , ns $\triangleq$ not significant).

**Extended Data Table 2.**
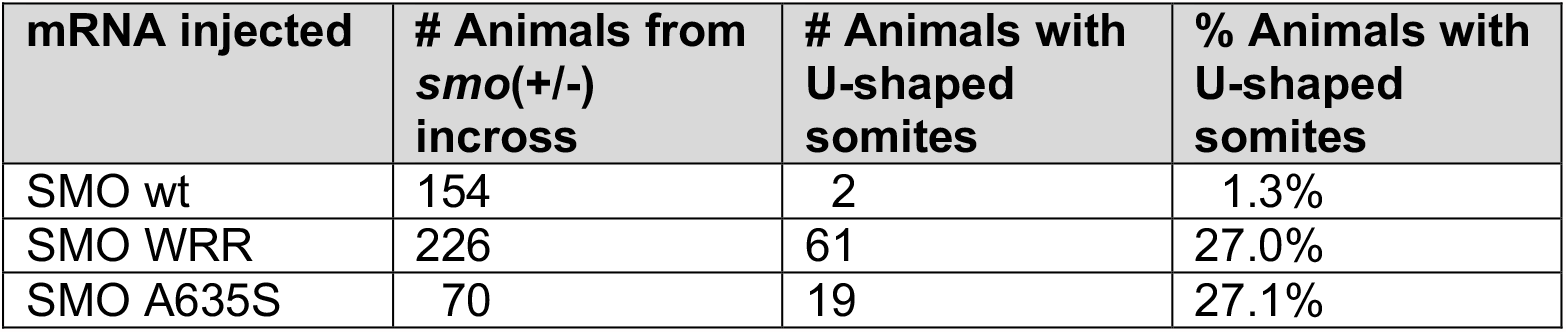
Quantification of phenotypes for zebrafish embryogenesis studies. The # and % of animals exhibiting U-shaped somites (indicative of a failure in Hh signaling during somitogenesis^55,98-100^) are indicated. Note that in both the WRR and A635S mutant conditions, close to 25% of animals exhibited U-shaped somites, consistent with the Mendelian inheritance of a null *smo* allele from the initial heterozygous incross.

| mRNA injected | # Animals from *smo*(+/-) incross | # Animals with U-shaped somites | % Animals with U-shaped somites |
| --- | --- | --- | --- |
| SMO wt | 154 | 2 | 1.3% |
| SMO WRR | 226 | 61 | 27.0% |
| SMO A635S | 70 | 19 | 27.1% |

